# Inbreeding preference emerges in female guppies in a socially complex setting

**DOI:** 10.1101/2025.03.20.644384

**Authors:** Amanda Viving, Léa Daupagne, David Wheatcroft, Niclas Kolm, John L. Fitzpatrick

**Author notes:** Contributing authors. These authors contributed equally to this work (Amanda Viving, Léa Daupagne).

## Abstract

Animals are typically expected to avoid mating with relatives due to the costs associated with incestuous matings. Yet for more than four decades, theoretical models have consistently suggested that animals should tolerate, or even prefer, mating with relatives under a broad range of conditions. However, empirical studies that evaluate inbreeding avoidance under alternative social and sexual contexts remain scarce. Here, we investigate how experimental variation in sexual and social complexity influence precopulatory inbreeding avoidance behaviors in the guppy *Poecilia reticulata*, a species known to experience inbreeding depression. In an integrated set of experiments, we examined if sexual and affiliative behaviors of virgin and experienced females and males were differentially directed towards either related or unrelated individuals. In simplistic social interactions, neither virgin nor experienced females or males showed a preference for related or unrelated partners. In contrast, we detected sex-specific responses to relatives in the more sexually and socially complex free-swimming arenas. Females exhibited a stronger preference towards related males regardless of mating experience, while male preference remained unchanged. Overall, these findings challenge previous reports of preference shifts between virgin and experienced female guppies and suggest that inbreeding avoidance behaviors may be less prevalent in complex social environments than previously thought.

## 1 Introduction

The question of why individuals mate with relatives when inbreeding has clear fitness costs (the so-called *inbreeding paradox*) has recently gained considerable theoretical and empirical interest (de Boer et al., 2021; Dorsey and Rosenthal, 2023; Montecinos et al., 2021; Reid et al., 2015; Townsend et al., 2019). Potential solutions to the inbreeding paradox have historically focused on the relative costs and benefits of inbreeding, particularly in terms of inclusive fitness. Specifically, inbreeding might be favored through kin selection, provided the potential negative effects of inbreeding are offset by the benefits associated with increasing the reproductive success of related individuals. Thus, choosing to mate with kin has been proposed as an adaptative strategy that enhances an individual’s indirect inclusive fitness (Kokko and Ots, 2006; Lehmann and Perrin, 2003; Szulkin et al., 2013; Duthie and Reid, 2016; Parker, 1979; Waser et al., 1986). Recently, Dorsey and Rosenthal (2023) offered a novel solution to the inbreeding paradox that focuses on the potential benefits of kin affiliation in sexual versus non-sexual situations. While sexual affiliation with relatives is often associated with fitness costs (i.e., through inbreeding depression), animals often benefit by interacting with relatives in non-sexual contexts. For instance, in species living in groups, kin preference can translate into fitness benefits by increasing individual growth (e.g., Thünken et al., 2016; Brown and Brown, 1996), potentially mitigating inbreeding costs. Since the benefits of non-sexual interactions with relatives require affiliative behaviors among kin, the evolution of mechanisms that reduce sexual interactions with kin may be constrained (Dorsey and Rosenthal, 2023), particularly when affiliative behaviors are selected at the adult phase. Yet, research on how sexual and non-sexual behaviors vary among social partners based on relatedness remains limited, as most studies focus on kin preferences during either sexual (e.g., mate choice) or non-sexual (e.g., juvenile and/or parental) life stages.

Inbreeding avoidance strategies can also be shaped by a range of factors, further complicating our ability to understand how preference for kin affiliation during non-sexual situation may constrain inbreeding avoidance during mate choice. For example, experimental approaches typically examine sexual behaviors without considering non-sexual behaviors and rarely examine inbreeding avoidance in complex social environments. Most studies employ a simple, dichotomous choice design, which may not fully capture the social dynamics found in natural systems (Carleial et al., 2020). Strong precopulatory mate preference may limited or be overridden through intra-sexual competition (Candolin, 2004; Forsgren, 1997; Kangas and Lindström, 2001; Sih et al., 2002) or sneak matings (Brockmann, 2001; Shuster and Wade, 2003). Indeed, previous studies show that an individual’s sexual and social behavior during free interactions in complex social settings may shift depending on the sex ratio, the number of surrounding conspecifics, and whether those conspecifics are related or not (Carazo et al., 2014; Daniel and Williamson, 2020; Martin and Long, 2015; Lymbery and Simmons, 2017). Additionally, studies exploring the effects of inbreeding avoidance within simple versus complex social environments usually do not account for key biological factors that may influence an individual’s propensity for inbreeding avoidance (de Boer et al., 2021). First, responses to inbreeding avoidance are predicted to be sex-specific, with females expected to exert stronger mate choice, while males are generally assumed to be less choosy (Blouin and Blouin, 1988; de Boer et al., 2021; Gerlach and Lysiak, 2006). Second, mating status may influence inbreeding avoidance, as virgin individuals are expected to mate indiscriminately to ensure reproductive assurance, while mated individuals are expected to be more selective, with choosiness increasing with mating experience (Kokko and Ots, 2006; Jennions and Petrie, 1997, 2000; Tanner et al., 2019). Indeed, evidence suggests that females tend to “trade-up” in genetic quality during sequential mate choice by mating with more compatible (e.g. Svensson et al., 2010; Bradley et al., 1990) or attractive males in later mating opportunities (e.g. Pitcher et al., 2003). In the context of inbreeding, this theory is supported by a recent study from Daniel and Rodd (2016) showing that virgin female guppies show no kin discrimination during their first mating episode, but avoid incest once mated. However, this evidence is challenged by two meta-analyses (de Boer et al., 2021; Richardson and Zuk, 2023), which indicate that mating status generally has limited influence on inbreeding avoidance and female choosiness. Nevertheless, these meta-analyses also highlight the general lack of studies considering mating status as a factor, emphasizing the need for its inclusion in future research (de Boer et al., 2021; Richardson and Zuk, 2023).

Here, we investigated how sex, mating status, and the complexity of the sexual-social environment influence behavioral affiliations for related versus unrelated individuals in both sexual and nonsexual contexts in the guppy (*Poecilia reticulata*). The guppy is an internally fertilizing, live-bearing freshwater fish, with a non-resource based, polygynandrous mating system. In the wild, guppies experience dry seasons that may create isolated small populations, where relatives are more likely to encounter one another, increasing the risk of inbreeding (Griffiths and Magurran, 1997). Inbreeding depression can be severe in guppies, as inbred offspring exhibit reduced survival rates (Nakadate et al., 2003; Oosterhout et al., 2007), lower courtship displays and sexual coloration (Van Oosterhout et al., 2003), and diminished sperm counts (Zajitschek and Brooks, 2010). Although females are generally the choosier sex, males may also display mating preferences, as they often encounter several females simultaneously (Houde, 1997*c*). Moreover, sperm production requires a refractory period between successive matings (Pilastro and Bisazza, 1999), prompting males to allocate sperm to the highest-quality female when reserves are limited (Dosen and Montgomerie, 2004; Andersson, 1994). Guppies express mating preference towards traits such as body size, female receptivity and male body coloration (Houde, 1997b; Corral-López et al., 2017; Auld et al., 2016), and can base their mating decision through kin recognition mechanisms using visual and/or olfactory cues (Penn and Frommen, 2010; Griffiths and Magurran, 1999). Kin recognition can be achieved through familiarity, where an individuals considers any conspecific that shares a degree of familiarity as kin, and phenotype matching, where an individual use a template of its own phenotype and/or that of its familiar kin to later evaluate others (Penn and Frommen, 2010). Given the risk of inbreeding in the wild, the associated fitness costs, and guppies’ ability to distinguish kin through various mechanisms, it is expected that guppies have evolved precopulatory inbreeding avoidance through mate choice (Daniel and Rodd, 2016). However, previous studies have shown mixed results (Viken et al., 2006; Guevara-Fiore et al., 2010; Pitcher et al., 2008; Zajitschek and Brooks, 2010; Daniel and Rodd, 2016). Moreover, guppies form stable social networks (Croft et al., 2003), which can promote preferential interactions among kin in non-sexual contexts. Such kin-based social bonds may provide cooperative benefits, such as increased survival or access to resources (e.g. Liévin-Bazin et al., 2019), potentially influencing the trade-off between affiliating with kin and avoiding inbreeding. If these non-sexual interactions are particularly beneficial among adult relatives, selection may act against strict precopulatory inbreeding avoidance in favor of maintaining social cohesion. By using individuals of both sexes with different mating statuses within environments of varying sexual-social complexity, this study aimed to identify factors that may constrain or modify the costs and benefits associated with inbreeding. We predicted that (1) females would exhibit stronger inbreeding avoidance than males, (2) virgins would be less choosy than experienced individuals, especially among females, (3) individuals would preferentially interact with unrelated individuals in a sexual context, while such preference would not be observed or reversed in a non-sexual context.

## 2 Methods

### 2.1 Experimental fish and rearing conditions

Fish used in this study were derived from Trinidadian guppy populations from high-predation areas of the Quare River. Stock populations were maintained in groups of several hundred fish, with regular movement of fish among stock populations. From these stock populations, we generated 24 distinct families of full sibling offspring, each generated through an independent cross between a single virgin female and male guppy. To further reduce the risk of inbreeding, 24 females and 24 males were randomly selected from two different stock tanks, with each pair formed by combining a female from one tank and male from the other. These pairs were then housed in 4-L tanks until they produced their first brood. The fry from each brood were housed in groups of no more than six, and were separated into sex specific tanks when males reached sexual maturity (at approximately *∼* 2-3 months). This method resulted in full siblings (r *∼* 0.5) and unrelated individuals (r *∼* 0). The tanks were visually isolated from the opposite sex, as male color patterns can influence future mate choice decisions in females (Breden et al., 1995; Hughes et al., 1999). The fish bred for this experiment were 9-11 months old at the time of the experiment. The fish were kept on a 12:12 light:dark cycle at 26-28°C and fed dry flakes and Artemia.

### 2.2 Experimental procedure

The role of relatedness in shaping sexual and social interactions among guppies was assessed in three separate and successive experiments (Fig. 1). Each focal individual was tested in each experimental setting. In Experiment 1 (dichotomous mate choice), a focal individual was presented with two size-matched stimuli individuals that differed in their relatedness (related versus unrelated, Fig. 1 A). In Experiment 2, four individuals from one family (two females, two males) were placed together with four individuals from another family (two females, two males) in a free-swimming arena and reproductive behaviors were recorded (Fig. 1 B). In Experiment 3, two same-sex individuals from one family were placed together with two same-sex individuals from three other families (i.e., forming groups of eight fish, Fig. 1 C). The experiments were divided equally into morning and afternoon sessions.

**Fig. 1:**
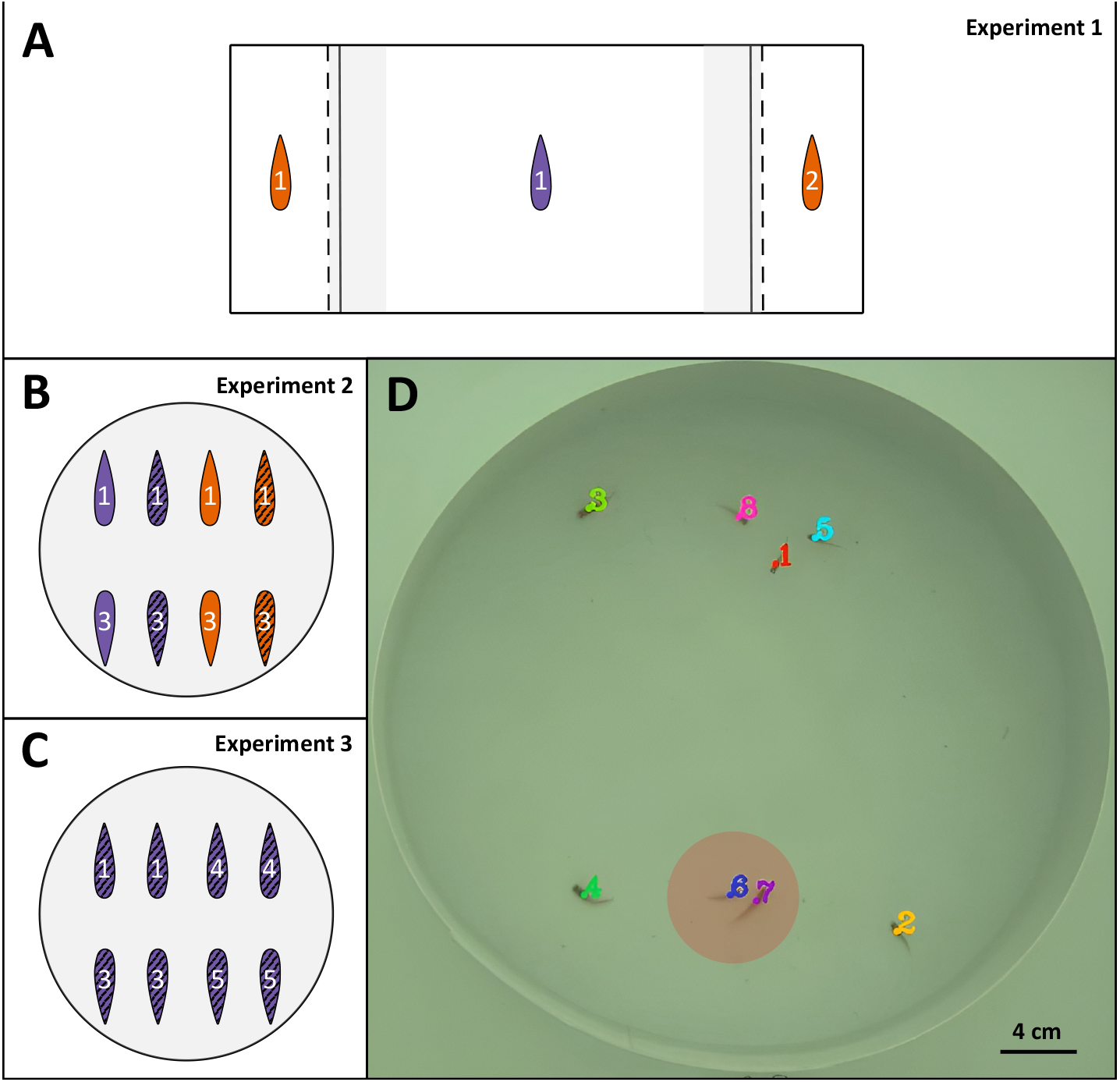
Experimental settings used in the study. In panels A, B, and C, females are shown in purple, males in orange, and the numbers correspond to families. Striped patterns indicate sexually experienced individuals, while plain (unstriped) patterns represent virgin individuals. **A**: Schematic representation of a dichotomous mate choice tank (Experiment 1). The focal individual in the main compartment (female or male, virgin or experienced) faced two stimulus individuals (related and unrelated) of the opposite-sex placed at random in the left and and right stimulus compartments. Two opaque barriers (solid black lines) prevented visual contact between the stimulus compartments and the chooser individual in the main compartment. After habituation, the opaque dividers were lifted and mate choice was assessed by recording the time the focal individual spent in either the left or right association zone (24 cm x 5 cm, here shown in grey areas). **B**: Schematic representation of a free-swimming mate choice arena (Experiment 2). Four individuals from a single family, representing both sexes and mating statuses, were introduced to interact with four individuals from a unrelated family of identical composition. Prior to this interaction, the two family groups had no previous exposure to one another. **C**: Schematic representation of a free-swimming social interactions arena (Experiment 3). Two same-sex individuals from a single family were introduced to interact with two individuals from each of three unrelated families. **D**: Example of a tracked video generated by idtracker.ai, i.e. an original input video overlayed with the individual trajectories and labels, enabling identification of individuals engaging in sexual behaviors. *idtracker*.*ai* provides the position of the centroid of each individual in pixel coordinates for avery frame. Individuals were considered associated when located within a 3.52 cm radius of each other (see Methods 2.2.3); e.g. individual 7 is within individual 6’s association zone (red area), indicating an association between the pair at that frame. Each focal individual tested in Experiment 1 was also tested in Experiment 2 and 3.

#### 2.2.1 Acclimation and mating status definition

All fish were virgins as the start of the experiment. Five days prior to Experiment 1, each individual was acclimated in a medium-sized (7-L) tank containing white gravel, along with a sibling of the same sex. This setup was designed to familiarize the fish with a white background similar to that used in the experimental trials. One day prior to Experiment 1, a focal fish (either male or female) was housed together in a 4-L tank with one individual from a different family of the same sex (same-sex housing) to ensure they remained virgin, or, of opposite-sex (opposite-sex housing) to provide individuals with mating experience. These pairings were between unrelated families with no prior experience, and each pair was kept together for approximately 24 hours to allow sufficient time for mating. Based on the protocol established by Daniel and Rodd (2016), mating typically occurs within this time period, although we did not directly observe mating. After the housing period, each fish was transferred to an individual 4-L tank enriched with plants and white gravel, where they remained for the duration of the experimental period.

#### 2.2.2 Experiment 1: dichotomous mate choice experiment

To test if mating status and sex influenced mate preferences, we first presented fish with a dichotomous choice of a related and unrelated potential partner. The dichotomous mate choice was conducted in 40 × 24 × 20 cm rectangular tanks divided into three compartments: one main compartment (27 × 24 × 20 cm) which housed the focal individual (female or male, virgin or experienced), and two smaller compartments (each 6 × 24 × 20 cm) which housed the two opposite-sex stimulus individuals, one of which was related to the focal individual and the other unrelated (see Fig. 1, A). The stimulus individuals were size-matched and the placement of the related and unrelated individuals in the smaller compartments was randomized. All tanks were filled to a depth of 5 cm. A transparent plastic barrier with small openings (allowing water flow between chambers) separated stimulus individuals from the focal individual, allowing the focal individual to use both olfactory and visual cues to assess stimulus individuals during the trials (Guevara-Fiore et al., 2009; Shohet and Watt, 2004; Houde, 1997*a,c*). Before each trial, the individuals were given a 30-min period to acclimatize to their environment. During this time, visual contact between the main and stimulus chambers was prevented by two opaque dividers. After habituation, the opaque barriers were removed and mate choice was assessed by recording the time the focal individual spent in each association zone, an area of 5 cm in front of the respective stimulus compartments (24 × 5 cm, see Fig. 1, A). The time individuals spend in an association zone is a commonly used measure of mating preference that is predictive of mate preference during sexual interactions (Dougherty, 2020). Trials were recorded using a camera (Point Grey Grasshopper 3 4.1 megapixel camera with Fujinon CF25HA-1 lens) directly placed 1.5 m above the experimental tanks. To remove olfactory cues between trials, each tank was drained and refilled with fresh water following each trial.

In total, 96 trials were performed, testing 96 individuals (48 males, 48 females) with different mating status (24 virgin males, 24 experienced males, 24 virgin females and 24 experienced females). In a few cases, the focal fish went under the opaque barrier during the acclimation period (n=7, three virgin males, two experienced males and two experienced females) so the recordings were excluded from the analysis. This resulted in a final sample size of 89 trials being included in the statistical analysis. From the recording, mate preference was assessed using an automated tracking software (EthoVision XT 17, Noldus Information Technology). Video analyses were performed by a single observer (A.V.) blind to both mating status and relatedness.

#### 2.2.3 Experiment 2: free-swimming mate choice

The free-swimming mate choice was assessed in cylinder arenas (36 cm in diameter), containing eight individuals, each of whom had been tested as a focal subject in the dichotomous mate choice experiment the previous day (see Fig. 1, B). In each trial, four individuals from a single family, representing both sex and mating statuses, were allowed to freely interact with four individuals from an unrelated family of identical composition. These two families had not experienced one another prior the trial. The individuals were acclimated in smaller isolated chambers (5 × 6 cm) within the cylinder arena for 15 minutes, which were then lifted and the behaviors of the individuals were recorded for 30 min.

Behavioral scoring was assessed using Behavioural Observation Research Interactive Software *BORIS* (Friard and Gamba, 2016). During observations, tracked videos generated by *idtracker*.*ai* (Romero-Ferrero et al., 2019) were simultaneously playing, allowing the identification of individuals engaging in sexual behaviors. In females, we scored three sexual behaviors : orienting behavior (turning and facing the displaying male), gliding behavior (moving in a smooth gliding fashion towards the male) and approaching behavior (moving towards the displaying male) (Houde, 1997*c*). In males, we scored the time spent pursuing the female (following the female, < 2 body lengths away), the number of sigmoid displays (quivering with an S-shaped arch of the body to the side or in front of the female), the number of successful matings, and the number of forced mating attempts (an alternative mating strategy where the male moves behind the female and thrusts his gonopodium at the female’s gonopore) (Houde, 1997*c*). In total, 12 trials were performed (each consisting of 8 individuals). Video analyses were conducted by a single observer (A.V.) blind to both mating status and relatedness.

We also examined sexual and social interactions within the arenas by recording the spatial proximity between each pair of individuals. Using identity-registered centroid trajectories produced by *idtracker*.*ai* version 5.2 11. (Romero-Ferrero et al., 2019), we calculated Euclidean distance matrices for each recording frame. We defined two individuals as being associated if they were located within a 3.52 cm radius, i.e. < 2 body lengths apart. This distance threshold was determined based on our data, where individuals measured an average of 1.76 cm, making two body lengths equal to 3.52 cm.

#### 2.2.4 Experiment 3: free-swimming social interactions

The following day, the same cylinder arenas (36 cm in diameter) were used to investigate social interactions in a non-sexual context (see Fig. 1, C). Here, two same-sex individuals from a single family were introduced to interact with two individuals from each of three unrelated families. The spatial proximity between each pair of individuals was assessed in the same way that was described in Experiment 2 (2.2.3). In total, 12 trials were performed (each consisting of 8 individuals). Video analyses were conducted by a single observer (A.V.) blind to both sex and relatedness.

#### 2.2.5 Measuring morphological traits

One day prior to experiment 1, each fish was photographed with a Canon EOS 800D and all measurements were gathered through *ImageJ* version 1.54. (https://imagej.net/ij/) for subsequent morphological analyses of attractive traits including body size and total area of orange coloration in males (Karino and Matsunaga, 2002; Herdman et al., 2004; Corral-López et al., 2017). The difference in body size between stimulus individuals was calculated, as well as the difference in total orange coloration between stimulus males. There was no significant difference in body size between related and unrelated stimulus individuals (Wilcoxon rank-sum test: W = 4413.5, p = 0.19). Similarly, total orange coloration did not differ between unrelated and unrelated males (Wilcoxon rank-sum test: W = 1170, p = 0.38). The average body length difference was 1.96 mm (± SD, 1.53 mm) in females and 1.06 mm (± 0.77 mm) in males. The average body area difference was 15.1 mm^2^ (± 13 mm^2^) in females and 6.2 mm^2^ (± 4.4 mm^2^) in males. Furthermore, males had an average difference in total area of orange coloration of 1.25 mm2 (± 1.1 mm^2^).

### 2.3 Statistical analysis

All statistical analyses were performed using *R* version 4.3.2. We constructed linear and generalized linear mixed effects models (LMMs and GLMMs) using the *lme4* (Bates et al., 2015) and *glmmTMB* (Brooks et al., 2017) packages.

#### 2.3.1 Experiment 1: dichotomous mate choice

Strength of preference (SOP) scores were used to estimate individual preference for either stimuli within each trial. SOP scores were calculated following the equation:

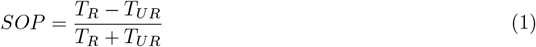

where *T*_*R*_ is the time spent in the related association zone and *T*_*UR*_ is the time spent in the unrelated association zone. This provides a SOP range from - 1 to 1. A SOP closer to - 1 indicates a preference towards unrelated individuals (i.e. inbreeding avoidance), a SOP closer to 1 indicates a preference towards related (i.e. kin preference) and a SOP of 0 indicated an unbiased mating preference.

To determine whether guppies, overall, preferred related or unrelated individuals, a one-sample t-test was performed on the SOP scores obtained from the total duration of the trial. SOP scores were then used as the response variable in an LMM, while sex and mating status were included as fixed factors. Random factor accounted for potential variations due to family origin. SOP scores obtained across multiple choice intervals were additionally calculated to investigate consistent patterns of focal preferences (Table A2). Specifically, we quantified informed choice by considering association times after the focal individual had visited the association zones of both stimuli individuals. One trial was removed from the informed choice analysis as the focal individual visited the second stimuli individual late into the video, resulting in a final sample size of 88 trials. Two additional LMMs were performed in each sex to investigate whether differences in attractive traits between related and unrelated stimuli individuals could have influenced focal SOP. In males, the model was fitted with the differences in body length and orange coloration together with mating status as fixed factors while in females, the model was performed with the difference in body length together with mating status as fixed factors. The assumptions of normality and homoscedasticity required for LMMs were checked for all models. Two-way interactions were initially included in all models, but were excluded from the final models as they were non-significant.

#### 2.3.2 Experiment 2: free-swimming mate choice

To examine whether virgin and experienced male and female guppies displayed a sexual preference for related or unrelated individuals in a free-swimming arena setting, GLMMs were performed. Three metrics of female mate choice (total count of orienting, approaching or gliding behaviors) and three metrics of male mate choice (total count of sigmoid display, pursuing and mating behaviors) were used as response variables in separate models. Focal mating status and relatedness were included as fixed effects, while arena identity was incorporated as a random effect to account for variability between arenas. Mating behaviors included both successful mating acts and forced mating attempts. Additionally, we constructed a single GLMM for each sex to evaluate the overall effect of relatedness and mating status on all sexual behaviors (Table 2, d and h). In these models, behavioral type was included as a random effect. This approach enabled us to estimate the influence of relatedness and mating status on the overall frequency of behaviors while accounting for variations in baseline counts across different behavioral types. We also examined whether the spatial proximity between all pairs or between female-male pairs only was influenced by their relatedness and respective mating status using a GLMM. As predictors had more than two categories, we used the *Anova* function from the *car* package to obtain significance effects. Significant main effects were then examined using pairwise Tukey post hoc comparisons in the *emmeans* package. Two-way interactions were initially included in all models, but were excluded from the final models as they were non-significant. A negative binomial distribution was used to account for overdispersion in the data.

#### 2.3.3 Experiment 3: free-swimming social interactions

To examine whether male and female guppies displayed a social preference for related or unrelated individuals in a non-sexual setting, a GLMM was performed with association duration as a response variable. Relatedness and sex were included as fixed effects and arena identity as a random effect to account for variability across arenas. A two-way interaction term was initially included, but was removed as non-significant. A negative binomial distribution was used to account for overdispersion in the data.

#### 2.3.4 Preference for relatives across experiments

To evaluate whether an individual’s preference for relatives remains consistent across experimental designs in a sexual context, we performed Spearman rank correlations between the strength of preference (SOP) for relatives in opposite-sex arenas (Experiment 2) and in dichotomous mate choice trials (Experiment 1). Additionally, we assessed the consistency of these preferences across both sexual and non-sexual contexts by performing Spearman rank correlations between (1) SOP for relatives in same-sex arenas (Experiment 3) and dichotomous mate choice trials (Experiment 1) and (2) SOP for relatives in opposite-sex arenas (Experiment 2) and same-sex arenas (Experiment 3). In Experiment 2, SOP was calculated following Equation 1 with *T*_*R*_ being the time spent in association with the two related individuals of the opposite-sex and *T*_*UR*_ being the time spent in association with the two unrelated individuals of the opposite-sex. In Experiment 3, *T*_*R*_ was the time spent in association with the related individual and *T*_*UR*_ was the time spent in association the six unrelated individuals (see Fig. 1 C).

##### Ethical statement

Animal housing and experimental protocols were performed in accordance with the guidelines of the animal research ethical board (permit number: 18352-2020).

## 3 Results

### 3.1 Experiment 1: dichotomous mate choice

During the 30-minute choice period, females spent an average of 68% of their time in either association zone (mean ± SD: 20.34 ± 3.60 min), while male guppies spent 58% (17.53 ± 4.02 min). On average, guppies did not show a significant overall preference for related or unrelated individuals (mean SOP = −0.016, *t(88)* = −0.47, *P-value* = 0.64). Neither sex nor mating status significantly influenced the strength of preference (SOP) for related partners (Table 1a, Fig. 2). However, related individuals were preferred when they were larger than unrelated individuals. Similar results were obtained when SOP scores were calculated at different choice intervals (Table A1, A2). When analyzed separately by sex, SOP for related individuals was also affected by the body size difference between stimuli (Table 1, b and c). In females, the SOP for related males was not influenced by differences in orange coloration between males (Table 1, b). SOP scores obtained across multiple choice intervals were additionally calculated to investigate consistent patterns of focal preferences

**Table 1:** Effect of sex and mating status on mate preference for related individuals.

| Response variable | N | Predictors | Estimates | SE | z | P-values |
| --- | --- | --- | --- | --- | --- | --- |
| (a) *SOP for relatives* | 89 | Body length | 0.05 | 0.02 | 3.23 | **0.002(\*\*)** |
|  |  | Sex (Male) | -0.06 | 0.06 | -1.13 | 0.26 |
|  |  | Mating status (Virgin) | -0.01 | 0.06 | -0.18 | 0.86 |
| (b) *Female SOP for related males* | 46 | Body length | 0.07 | 0.03 | 1.96 | **0.05(\*)** |
|  |  | Orange coloration | 0.02 | 0.02 | 0.94 | 0.35 |
|  |  | Mating status (Virgin) | -0.05 | 0.06 | -0.81 | 0.43 |
| (c) *Male SOP for related females* | 43 | Body length | 0.05 | 0.02 | 2.12 | **0.04(\*)** |
|  |  | Mating status (Virgin) | 0.06 | 0.08 | 0.78 | 0.45 |

Outputs of linear mixed models (LMMs) investigating (a) the effect of difference in body length, sex and mating status on strength of preference (SOP) for related individuals, (b) the effect of difference in body length, difference in orange coloration and mating status on female SOP for related males, and (c) the effect of difference in body length and mating status on male SOP for related females. For each sex, body traits were compared by subtracting the trait value of the unrelated stimuli from that of the related stimuli. Positive differences indicate larger values in the related stimuli, while negative differences indicate larger values in the unrelated stimuli. The total number of trials performed (N) is presented for each model, along with the standard error (SE), test statistic (z) and P-value for each effect. Significant P-values are shown in bold.

**Fig. 2:**
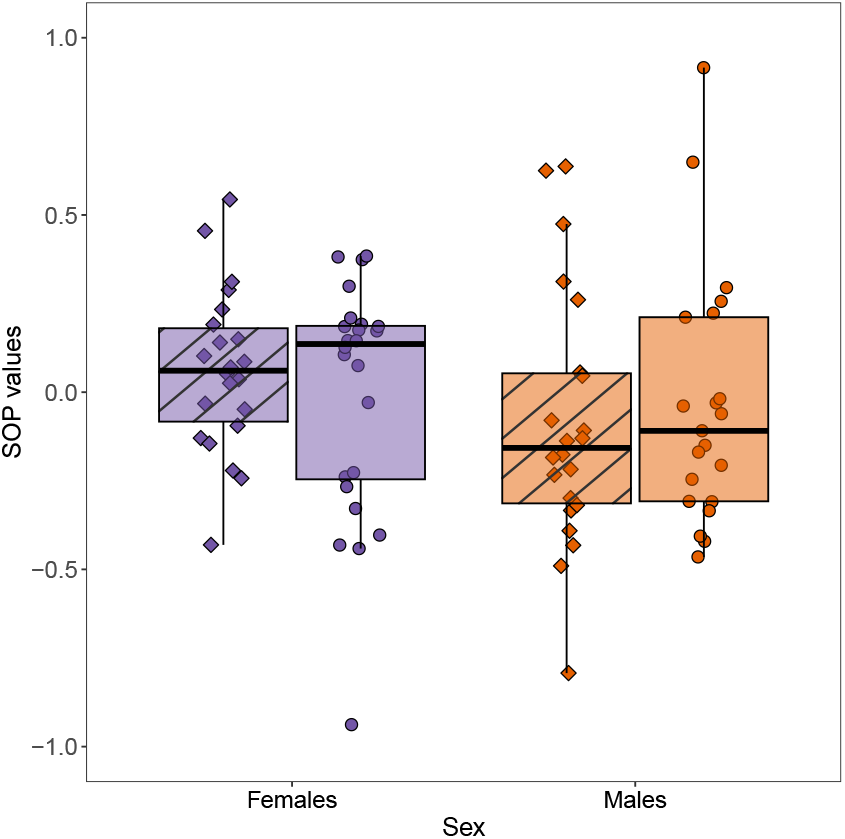
Strength of preference (SOP) scores obtained in females (purple) and males (orange) throughout the duration of a trial (0-30 minutes). Data are compared between experienced (striped) and virgin (plain) individuals in each sex. In the box plots, the horizontal lines indicate the median (50th percentile) and quartiles (25th and 75th percentiles) and the vertical whiskers indicate the range of the jittered data points.

### 3.2 Experiment 2: free-swimming mate choice

#### 3.2.1 Female behaviors

Each of the female behaviors investigated (gliding, orienting, and approaching behaviors) did not differ significantly depending on the relatedness to the focal male (GLMMs; Table 2, a, b and c, Fig. 3). However, when all sexual behaviors were analyzed, female responses to male courtship differed depending on relatedness, with females directing overall more responses towards males that were related (GLMM; Table 2, d). Additionally, virgin females directed significantly more gliding behaviors toward males than experienced females (GLMM; Table 2, b).

**Table 2:** Effect of relatedness and mating status on female and male sexual behaviors and association duration in mixed-sex arenas.

| Response variable | N | Predictors | Estimates | SE | *z* | *P-values* |
| --- | --- | --- | --- | --- | --- | --- |
| Female behaviors |  |  |  |  |  |  |
| (a) *Gliding behavior* | 46 | Relatedness (Unrelated) | -0.55 | 0.46 | -1.20 | 0.23 |
|  |  | Mating status (Virgin) | 1.33 | 0.50 | 2.68 | **0.007(\*\*)** |
| (b) *Orienting behavior* | 46 | Relatedness (Unrelated) | -0.21 | 0.15 | -1.42 | 0.16 |
|  |  | Mating status (Virgin) | 0.08 | 0.15 | 0.56 | 0.57 |
| (c) *Approaching behavior* | 46 | Relatedness (Unrelated) | -0.26 | 0.17 | -1.54 | 0.12 |
|  |  | Mating status (Virgin) | 0.01 | 0.17 | 0.08 | 0.94 |
| (d) **All sexual behaviors** | 46 | Relatedness (Unrelated) | -0.25 | 0.11 | -0.18 | **0.03(\*)** |
|  |  | Mating status (Virgin) | 0.17 | 0.11 | 1.52 | 0.13 |
| Male behaviors |  |  |  |  |  |  |
| (e) *Sigmoid display* | 46 | Relatedness (Unrelated) | -0.15 | 0.18 | -0.86 | 0.39 |
|  |  | Mating status (Virgin) | 0.14 | 0.18 | 0.78 | 0.44 |
| (f) *Pursuing behavior* | 46 | Relatedness (Unrelated) | 0.04 | 0.09 | 0.46 | 0.63 |
|  |  | Mating status (Virgin) | -0.22 | 0.09 | -2.45 | **0.01(\*\*)** |
| (g) *Mating behavior* | 46 | Relatedness (Unrelated) | 0.02 | 0.23 | 0.10 | 0.92 |
|  |  | Mating status (Virgin) | -0.61 | 0.23 | -2.63 | **0.008(\*\*)** |
| (h) **All sexual behaviors** | 46 | Relatedness (Unrelated) | -0.06 | 0.11 | -0.50 | 0.61 |
|  |  | Mating status (Virgin) | -0.17 | 0.11 | -1.53 | 0.12 |
| Association duration | | | | | $\chi^2$ | |
| (i) *Female-male pairs* | 192 | Relatedness |  |  | 0.66 | 0.42 |
|  |  | Mating status |  |  | 11.59 | **0.003(\*\*)** |
| (j) *All pairs* | 336 | Relatedness |  |  | 0.25 | 0.61 |
|  |  | Mating status |  |  | 18.41 | **<0.001(\*\*\*)** |
|  |  | Sex |  |  | 20.42 | **<0.001(\*\*\*)** |

After testing for an effect of relatedness and mating status on each female (a, b, and c) and male (e, f, and g) sexual behaviors in 12 mixed-sex arenas using univariate generalized linear mixed models (GLMMs), we assessed the overall effect of relatedness and mating status by pooling all sexual behaviors into a single GLMM with behavioral type included as a random effect (d and h). For models a-h, we report the $z$ -statistic, obtained from the `summary()` function in R. The $z$ -statistic is computed as the estimated coefficient divided by its standard error (SE) and assesses the significance of each fixed effect in the model. For GLMMs on association durations (i and j), we used the `Anova` function from the `car` package to obtain significance effects as predictors have more than two categories and reported the $\chi^2$ test statistic from likelihood ratio tests. Significant effects were further examined using pairwise Tukey post hoc comparisons in the `emmeans` package, as described in the main text. The total number of individuals or pairs assessed (N) is presented for each model, along with the SE, test statistic ( $z$ or $\chi^2$ value) and $P$ -value for each effect. Significant $P$ -values are shown in bold.

**Fig. 3:**
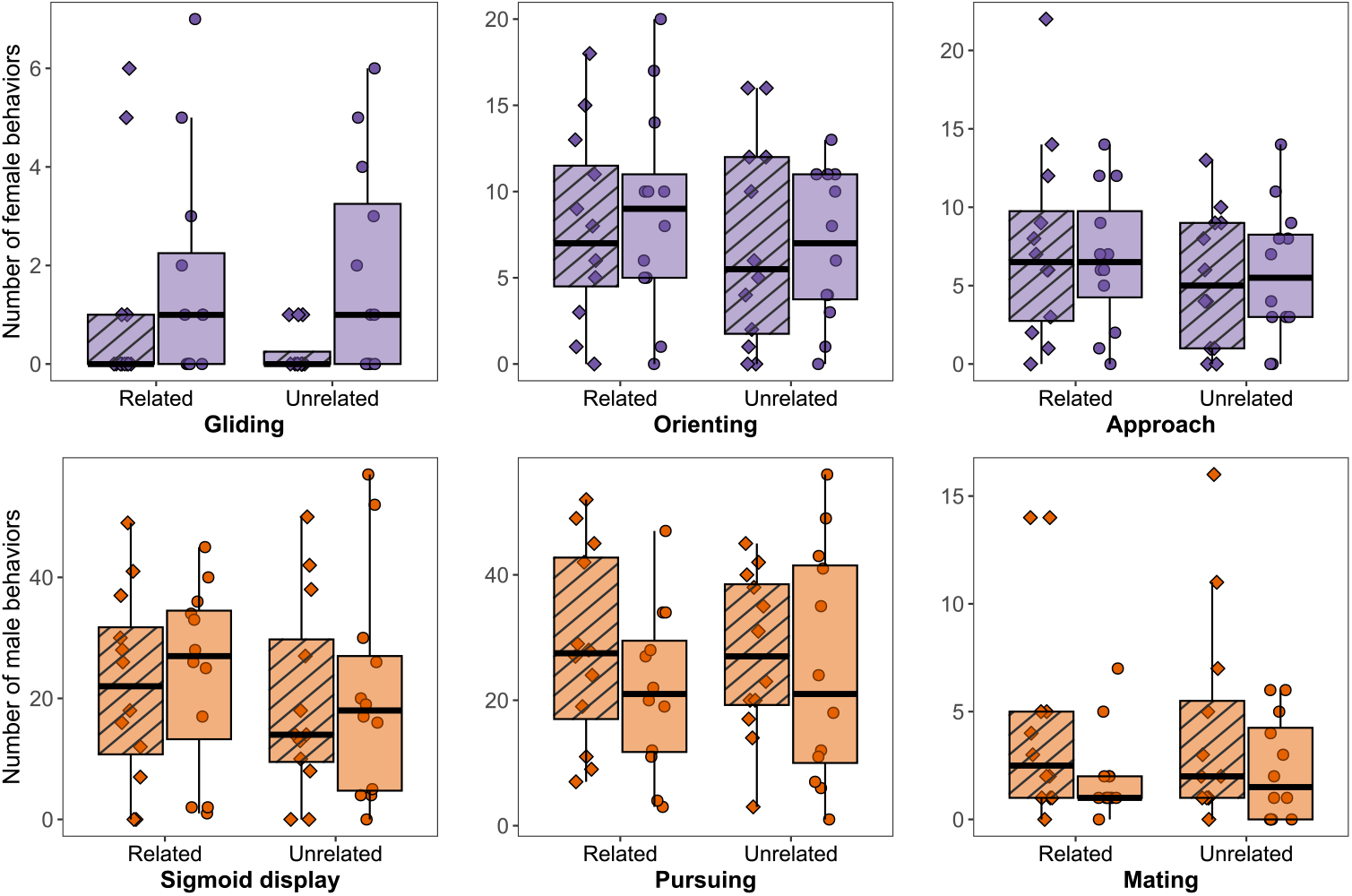
Total number of sexual behaviors observed in females (gliding, orienting and approaching behaviors) and males (sigmoid displays, pursing and mating behaviors), directed toward related and unrelated individuals in each arena for 30 minutes. Data are compared between experienced (striped) or virgin (plain) individuals in each sex (females = purple and males = orange). In the box plots, the horizontal lines indicate the median (50th percentile) and quartiles (25th and 75th percentiles) and the vertical whiskers indicate the range of the jittered data points.

#### 3.2.2 Male behaviors

Similarly to females, none of the male behaviors investigated separately (sigmoid displays, pursuing and mating behaviors), significantly differed depending on relatedness to the focal male (GLMMs; Table 2, e, f and g, Fig. 3). However, virgin males performed significantly fewer pursuing and mating behaviors than experienced males (GLMMs; Table 2, f and g, Fig. 3). When all sexual behaviors were analyzed, males did not significantly bias their behaviors depending on relatedness or mating status (GLMM; Table 2, h).

#### 3.2.3 Association duration

The frequency of female-male pair associations did not significantly differ based on the relatedness of the individuals involved (Table 2, i). However, mating status had a significant effect on the frequency of female-male pair associations (Table 2, i). Specifically, virgin individuals associated less frequently compared to pairs of experienced individuals (*z* = −2.98, *P-value* = 0.008) or virgin-experienced individuals (*z* = −3.06, *P-value* = 0.006).

The same pattern was observed when investigating the frequency of all pair associations (Table 2, j). The relatedness did not influence the frequency of associations (Table 2, j), but virgin individuals appeared to associate less frequently compared to pairs of experienced individuals (*z* = −3.76, *P-value* < 0.001) or virgin-experienced individuals (*z* = −3.89, *P-value* < 0.001). The sex of the individuals involved also significantly influenced the association frequency (Table 2, j). Male-male pairs associated significantly less than both male-female pairs (*z* = −4.25, *P-value* < 0.001) or female-female pairs (*z* = −3.73, *P-value* < 0.001).

### 3.3 Experiment 3: free-swimming social interactions

The frequency of pair associations did not significantly differ based on the relatedness of the individuals involved (Table 3, Fig. 4). However, males associated significantly less frequently compared to female pairs (Table 3, Fig. 4).

**Table 3:** Effect of relatedness and sex on association duration in same-sex arenas.

| Response variable | N | Predictors | Estimates | SE | *z* | *P-values* |
| --- | --- | --- | --- | --- | --- | --- |
| Association duration | 336 | Relatedness (Unrelated) | -0.05 | 0.08 | -0.66 | 0.51 |
|  |  | Sex (Male) | -1.65 | 0.27 | -6.09 | **<0.001(\*\*\*)** |

We evaluated the effect of relatedness and sex on association duration in 12 same-sex arenas using a GLMM with arena type included as a random effect. The total number of pairs assessed (N) is presented along with the standard error (SE), test statistic (*z*) and *P-value* for each effect. Significant *P-values* are shown in bold.

**Fig. 4:**
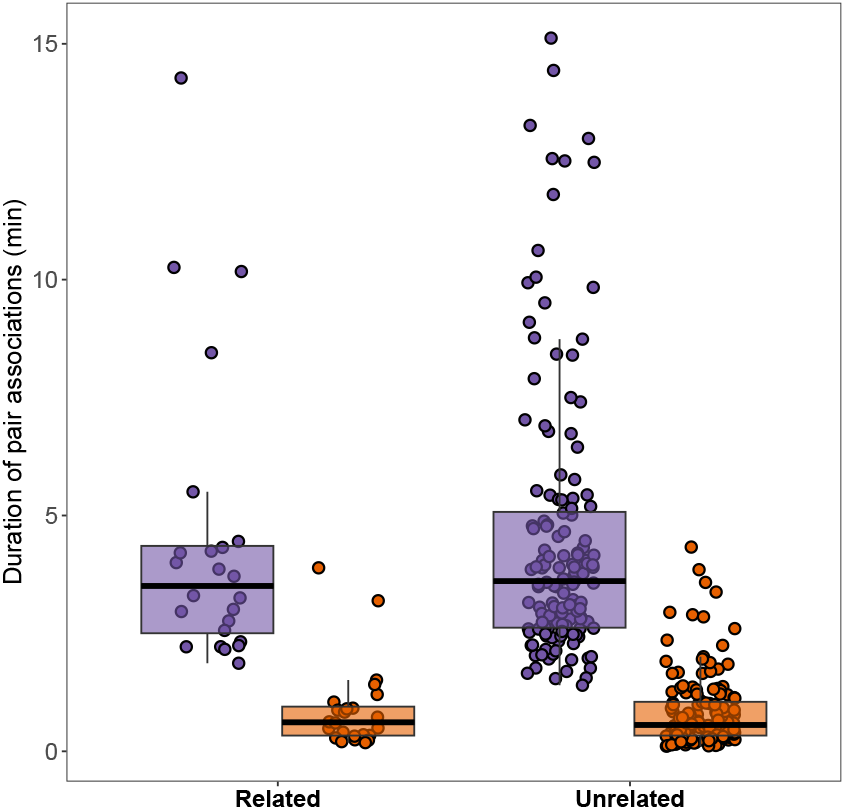
Association duration (min) between related and non-related pairs in same-sex arenas (females = purple, males = orange) analysed for 30 minutes. In the box plots, the horizontal lines indicate the median (50th percentile) and quartiles (25th and 75th percentiles) and the vertical whiskers indicate the range of the jittered data points.

#### 3.3.1 Preference for relatives across experiments

In both males and females, no significant correlation was observed between the strength of preference (SOP) for opposite-sex relatives in free-swimming arenas and in dichotomous mate choice trials (females: *r*_46_ = 0.005, P = 0.97; males: *r*_43_ = 0.17, P = 0.28; Fig. 5 A), indicating an individual’s preference for relatives is not consistent across experimental designs in a sexual context. Similarly, SOP for relatives in same-sex arenas did not correlate significantly with SOP in dichotomous mate choice trials (females: *r*_46_ = 0.24, P = 0.11; males: *r*_43_ = −0.08, P = 0.60; Fig. 5 B) or in opposite-sex arenas (females: *r*_46_ = −0.14, P = 0.36; males: *r*_43_ = 0.11, P = 0.49; Fig. 5 C), suggesting that an individual’s preference for relatives is not consistent across sexual and nonsexual contexts.

**Fig. 5:**
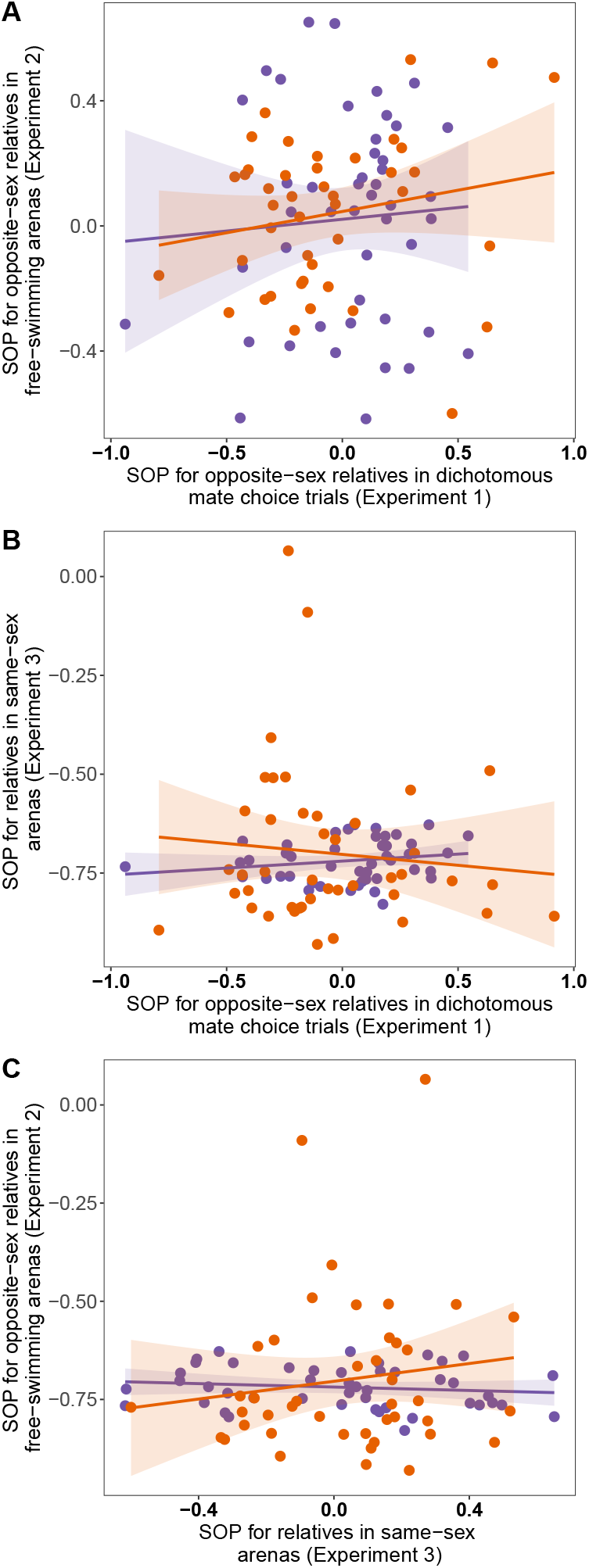
Pairwise relationship between (A) strength of preference (SOP) for opposite-sex relatives in free-swimming arenas (Experiment 2) and SOP for opposite-sex relatives in dichotomous mate choice trials (Experiment 1), (B) SOP for relatives in same-sex arenas (Experiment 3) and SOP for opposite-sex relatives in dichotomous mate choice trials (Experiment 1), (C) SOP for opposite-sex relatives in free-swimming arenas (Experiment 2) and SOP for relatives in same-sex arenas (Experiment 3). Data points represent individual metrics (females = purple, males = orange). Full lines depict linear regression lines fitted separately for each sex, and shaded areas represent their corresponding confidence intervals

## 4 Discussion

Contrary to longstanding expectations, we found little evidence that guppies avoid mating with relatives regardless of the experimental conditions. Moreover, despite differences in sexual behaviors between virgin and mated females and males, we found no evidence that mating status influenced mate preference for related versus unrelated mates in the guppy. However, female preference for related males emerged in environments with higher social complexity. In free-swimming trials, females were more responsive to male courtship from related males, suggesting that social context plays a role in shaping female mating preferences. Despite the absence of inbreeding avoidance, female and male guppies showed a clear preference for body size in dichotomous mate choice trials, suggesting body size may be a more salient determinant of mate choice. Furthermore, the overall absence of inbreeding avoidance observed in our study is not explained by non-sexual kin affiliation, as same-sex pair associations were not influenced by relatedness.

Regardless of their mating status, female guppies displayed an overall preference for related males in the complex social experimental setup when all female sexual behaviors were analyzed collectively. The costs associated with inbreeding depression are expected to generate selection for inbreeding avoidance (Pusey and Wolf, 1996; Blouin and Blouin, 1988). However, avoiding inbreeding may not always be advantageous, particularly if inbreeding costs are offset by indirect benefits of mating with relatives via kin selection. Kin preference through mate choice has been mainly documented in species that do not experience inbreeding depression such as the African cichlid fish (*Pelvicachromis taeniatus*) (Thünken et al., 2007), the ground tit (*Parus humilis*) (Wang and Lu, 2011) and the white’s skink (*Liopholis whitii*) (Bordogna et al., 2016). Interestingly, biased mating preference towards kin has also been found in species suffering from inbreeding depression including the American crow (Townsend et al., 2019) and the fruit fly (*Drosophila melanogaster*) (Loyau et al., 2012; Ehiobu et al., 1989). However, guppies are known to suffer from inbreeding depression (Nakadate et al., 2003; Oosterhout et al., 2007; Van Oosterhout et al., 2003; Zajitschek and Brooks, 2010) and we found no variation in association duration among female-male pairs based on relatedness. Whether preferential female association with related males could be explained by females gaining inclusive fitness benefits (Parker, 1979), remains an interesting avenue for future investigations.

In virgin females, our results align with previous mate choice experiments that reported no female preference for unrelated males through phenotype matching using either dichotomous choice trials (Viken et al., 2006; Zajitschek and Brooks, 2010; Guevara-Fiore et al., 2010) or free-swimming arenas (Pitcher et al., 2008; Daniel and Rodd, 2016). This supports theoretical models predicting that virgin females should be less choosy and/or more responsive to male courtship as rejecting a mating opportunity carries the risk of dying unmated (Jennions and Petrie, 2000; Kokko and Ots, 2006). Interestingly, we did not find evidence of a mate preference shift toward unrelated males when experienced females were used, in either the free-swimming or the dichotomous mate choice experiment. This suggests that for inbreeding avoidance female guppies do not alter their choosiness based on their current mating status. This finding contrasts with the results of Daniel and Rodd (2016), where the authors found that experienced females preferred unrelated males, while virgins did not display any preference. Although theory suggests that experienced females should choose males based on traits that indicate higher quality, such as genetic relatedness, our results match with two recent meta-analyses that found no evidence that virgin females are less choosy than mated females (Richardson and Zuk, 2023) or that female mating status influences inbreeding avoidance (de Boer et al., 2021), respectively. While Daniel and Rodd (2016) study relied on live behavioral observations, we used video recordings and tracking software to account for individual identities, potentially providing more accurate quantifications of female mate preferences.

Male guppies, irrespective of their mating status, exhibited no consistent preference based on relatedness in either experimental setup. The overall absence of strong sexual preferences in males aligns with theoretical expectations that, as the mate-limited sex, males prioritize reproductive opportunities over mate selectivity (Dosen and Montgomerie, 2004). Theoretical models (Kokko and Ots, 2006; Parker, 1979, 2006; Puurtinen, 2011; Waser et al., 1986) emphasize that inbreeding tolerance might be adaptive in males even when inbreeding depression occurs due to the opportunity cost in terms of missed opportunities to outbreed. This is particularly true for poeciliid fishes, where reproductive success is strongly tied to the number of mates acquired (Blouin and Blouin, 1988). Moreover, the evolution of male inbreeding avoidance behaviors in guppies may mainly be driven by female mate choice, as it been shown in polygynous primates (Tennenhouse, 2014). Here, we did not detect female mate preference towards unrelated males which additionally limits the selective pressures that might favor male inbreeding avoidance in guppies.

The benefits of non-sexual interactions among kin may limit the evolution of precopulatory inbreeding avoidance (Dorsey and Rosenthal, 2023). In salmonids (*Salmo salar* and *Oncorhynchus mykiss*), juveniles in shoals of related individuals grow faster and display fewer antagonistic behaviors than do shoals of unrelated individuals (Brown and Brown, 1996). Similarly, growth was higher in kin-only shoals of juvenile *Pelvicachromis taeniatus* than in mixed groups, indicating fitness benefits of kin shoaling. In guppies, Hain and Neff (2007) found that juveniles preferred to shoal with half-siblings than with unrelated individuals in the laboratory, while significant kin structure was found by Piyapong et al. (2011) in wild juvenile guppy shoals, although only in high predation environments. Guppies may therefore gain potential benefit from shoaling preferentially with kin at the juvenile stage. Here, we found no kin structure in pair associations within same-sex arenas, suggesting that non-sexual cooperation among adult kin is not a mechanism underlying the general absence of precopulatory inbreeding avoidance seen in this study. This may be explained by the reduced occurrence of shoaling behaviors performed by adults in natural populations. Although shoaling is a common defense against fish predation at the adult phase (Pitcher, 1993; Croft et al., 2006), previous studies consistently showed no evidence of kin structure in adult guppy aggregations of populations subjected to high predation pressure (Hain and Neff, 2007; Russell et al., 2004) where shoaling propensities should be maximized.

Precopulatory inbreeding avoidance mechanisms may also not evolve in guppies if they depend on sperm competition and cryptic female choice to mitigate the risks of inbreeding. The potential for such postcopulatory avoidance mechanisms is high in guppies as they display a highly polygynandrous mating system (Evans and Magurran, 1999), with females mating with several males each time they are receptive and males engaging in forced copulations (Magurran and Seghers, 1994; Pilastro and Bisazza, 1999). Moreover, females can store sperm for several months (Houde, 1997*c*; Liley, 1966), and guppies exhibit last male sperm precedence (Evans and Magurran, 2001). There is some evidence that postcopulatory avoidance mechanisms could act as a filter against incompatible sperm and minimize fertilization by sperm from mating partners with overly similar genotypes (Fitzpatrick and Evans, 2014; Gasparini and Pilastro, 2011), although counter examples show no bias fertilization toward less closely related males during competitive fertilizations (Evans et al., 2008; Pitcher et al., 2008). The overall prevalence of both pre- and postcopulatory inbreeding avoidance mechanisms may also depend on the severity of inbreeding depression that may be different between wild populations or laboratory raised populations. In addition, males often migrate in their natural environment (Croft et al., 2003), lowering the chance of meeting a relative to mate with, which may also reduce the propensity of inbreeding avoidance in guppies. Although our fish are laboratory raised, the expression of inbreeding avoidance before mating might still be low due to conserved selective pressures.

In conclusion, our findings highlight a limited role for precopulatory mechanisms in inbreeding avoidance among guppies. Female preferences for related mates in certain contexts may reflect kin selection benefits, while males appear to prioritize reproductive opportunities over relatedness. Furthermore, the absence of kin-structured associations in same-sex arenas indicates that examining nonsexual interactions offer little insight into resolving the inbreeding paradox. Together, these results indicate that dispersal and postcopulatory processes may be more important than premating selection in mitigating the costs of inbreeding depression in guppies.

## Authors’ contributions

A.V.: methodology, data curation, statistical analysis, writing—original draft, writing—review and editing; L.D.: methodology, data curation, statistical analysis, writing—original draft, writing—review and editing; N. K. and D.W.: conceptualisation, editing: J.L.F.: conceptualisation, funding acquisition, methodology, supervision, writing—review and editing. All authors gave final approval for publication and agreed to be held accountable for the work performed therein.

## Acknowledgments

We thank all current and past members of Fitzpatrick Lab, especially Mirjam Amcoff, for their support in fish feeding and monitoring. Experimental work used resources from Niclas Kolm laboratory. This work was funded by a Swedish Research Council Grant (2021-04615) to J.L.F. and a Carl Tryggers Foundation postdoctoral stipend (CTS 23:2794) to L.D.

## Data accessibility

Data and scripts will be available on Dryad.

## Conflict of interest declaration

We declare we have no competing interests.

## Appendix A

### A.1 Experiment 1: dichotomous mate choice

**Table A1:** The effect of difference in body length, sex and mating status on the strengh of preference (SOP) for related individuals when considering multiple time intervals.

| Response variable | N | Predictors | $\chi^2$ | *P-values* |
| --- | --- | --- | --- | --- |
| *SOP for relatives* | 89 | Body length | 14.16 | **<0.001(\*\*\*)** |
|  |  | Sex | 2.01 | 0.16 |
|  |  | Mating status | 0.02 | 0.89 |
|  |  | Time interval | 3.30 | 0.19 |

Outputs of a linear mixed models (LMM) investigating the effect of difference in body length, sex, and mating status on strength of preference (SOP) for related individuals when considering multiple time intervals. For each sex, body traits were compared by subtracting the trait value of the unrelated stimuli from that of the related stimuli. SOP scores comparisons were done across multiple time intervals: first 10 min, 10-20 min, and 20-30 min. The total number of trials performed (N) is presented, along with the test statistic ( $\chi^2$ ) and *P-value* for each effect. Significant *P-values* are shown in bold.

**Table A2:** The effect of sex and mating status on the strengh of preference (SOP) for related individuals accross multiple time intervals.

| Intervals | N | Predictors | Estimates | SE | *z* | *P-values* |
| --- | --- | --- | --- | --- | --- | --- |
| *0-10 minutes* | *89* | Sex (Male) | -0.10 | 0.08 | -1.12 | 0.26 |
|  |  | Mating status (Virgin) | 0.001 | 0.09 | 0.02 | 0.99 |
| *10-20 minutes* | *89* | Sex (Male) | -0.11 | 0.09 | -1.29 | 0.20 |
|  |  | Mating status (Virgin) | -0.01 | 0.09 | -0.16 | 0.87 |
| *0-20 minutes* | *89* | Sex (Male) | -0.11 | 0.07 | -1.64 | 0.11 |
|  |  | Mating status (Virgin) | 0.001 | 0.07 | 0.01 | 0.99 |
| *20-30 minutes* | *89* | Sex (Male) | -0.07 | 0.11 | -0.63 | 0.53 |
|  |  | Mating status (Virgin) | -0.01 | 0.11 | -0.01 | 0.99 |
| *0-end (IC)* | *88* | Sex (Male) | -0.08 | 0.06 | -1.32 | 0.19 |
|  |  | Mating status (Virgin) | -0.002 | 0.06 | -0.03 | 0.97 |
| *0-10 minutes (IC)* | *88* | Sex (Male) | -0.17 | 0.08 | -2.18 | **0.03(\*)** |
|  |  | Mating status (Virgin) | 0.01 | 0.08 | 0.13 | 0.90 |
| *10-20 minutes (IC)* | *88* | Sex (Male) | -0.05 | 0.09 | -0.57 | 0.57 |
|  |  | Mating status (Virgin) | -0.03 | 0.09 | -0.36 | 0.72 |

Outputs of a linear mixed models (LMM) investigating the effect of difference in body length, sex, and mating status on strength of preference (SOP) for related individuals. For each sex, body traits were compared by subtracting the trait value of the unrelated stimuli from that of the related stimuli. SOP scores comparisons were done across multiple time intervals: first 10 min, 10-20 min, first 20 min, 20-30 min, overall time after informed choice (IC), first 10 min after IC, 10-20 after IC. The total number of trials performed (N) is presented for each model, along with the standard error (SE), test statistic (*z*) and *P-value* for each effect. Significant *P-values* are shown in bold.

## Notes

### Competing Interest Statement

The authors have declared no competing interest.

